# A frameshift mutation in *PRICKLE1* causes skeletal dysplasia in Cheviot sheep

**DOI:** 10.64898/2026.10.07.757262

**Authors:** Oswald Matika, Megan G. Davey, Jorge del-Pozo, Heather A. Finlayson, Colin Farquharson, Denis J. Headon, James W. Kijas, Lynn McTeir, Cameron Batho-Samblas, Elizabeth Watson, Valentina Riggio, Jeffrey J. Schoenebeck, Tobias Schwarz, Katherine Ann Staines, Ricardo Pong-Wong, Zen H. Lu, Stephen C. Bishop, John A. Woolliams, Alan L. Archibald

## Abstract

Skeletal dysplasias can occur in animals, including humans, and are attributed to both environmental and genetic causes. As our understanding of the distinguishing phenotypes of different skeletal dysplasias improves, alongside elucidation of the mechanisms and genetic networks that contribute to skeletal development, identifying and controlling genetic mutations in livestock populations has become increasingly possible, including approaches beyond selective breeding alone. Within Scotland an ‘achondroplasia’ syndrome that was first described in the South Country Cheviot breed of sheep (OMIA 000004-9940), has continued to occur periodically within the same geographical locations, despite culling of affected lambs over many decades. Affected lambs exhibit rounded heads, short legs, a loss of hooves, abnormal stance and short twisted tails. As the condition has been observed in flocks and as progeny from specific ram and ewe matings, and cases have been co-twinned to normal siblings, this suggests a syndrome caused by a recessive genetic condition rather than environmental factors. Through homozygosity mapping and whole genome sequence analyses, we identified a 10 bp deletion in the open reading frame of the *PRICKLE1* gene that encodes prickle planar cell polarity protein 1 and is essential for normal vertebrate embryonic development. We identified additional phenotypes present in affected lambs, including a loss of phalanges, neural tube defects, vertebral and articular chondrocyte defects, hair orientation differences and polycystic kidneys, consistent with a loss of PRICKLE1 function based on comparisons to mouse mutant lines. These findings enabled the development of a simple DNA test for the identified *PRICKLE1* mutation that was made commercially available to facilitate the elimination of this genetic disease from sheep flocks.

## Introduction

Chondrodysplasias are developmental diseases affecting cartilage that lead to abnormal formation of the skeletal structures in vertebrates and include many of the most common disorders in humans and livestock species. The most common skeletal dysplasia in humans, achondroplasia, is caused by activating mutations in *FGFR3*^1^, resulting in shortening of the long bones of the limb, leading to shortened stature as well as complex malformations of the head, thorax and vertebral column. Loss of function mutations in *FGFR3*, meanwhile, cause elongating dysplasia of the long bones in both humans and sheep^2,3^, indicative of the conservation of developmental processes between livestock animals, model animals such as mice, and humans. As well as the FGF pathway, the genetic causes of many other chondrodysplasias have been identified within well characterised signalling pathways and transcription factors that are essential for cartilage formation and patterning during embryonic development in humans, companion and farmed animals, including mutations in the *BMPR1B, IHH, PTHRP*, *RUNX2, SHOX, SOX9*, and *WNT5A* genes^1,4,5,6^, as well as mutations in structural proteins essential for the integrity of cartilage and bone such as *ACAN, FBN1* and *COL2A1*^7^, and proteins associated with the mineralisation of cartilage and bone^8^.

While many phenotypes between different chondrodysplasias overlap, there are distinctions that characterise specific syndromes, including the type and locations of cartilage affected and the involvement of organs other than the skeleton^9^. In the majority of skeletal dysplasias such as achondroplasias, that are caused by mutations in *FGFR3,* limb shortening primarily impacts the proximal limb bones (rhizomelic). This is a result of narrow and distorted growth plates that lead to precocious ossification and dwarfism^10^. Skeletal dysplasias associated with human Robinow syndrome, that occur due to one of several mutations in components of the Planar Cell Polarity (PCP) pathway, including *WNT5A*, *DVL1* and *ROR2*, cause shortening of distal bones of the limb missing distal phalanges, nails or claws and disorganisation of the chondrocytes within the growth plate^11^ (reviewed^12^). Moreover, as the highly conserved PCP pathway provides the co-ordination of polarised cell behaviour such as directional growth, migration and patterning of many different organ systems, these skeletal dysplasias are frequently syndromic, affecting the nervous system, heart, dentition and genitals (reviewed^13^).

Charles Darwin first described chondrodysplasia in short legged Ancon sheep in his seminal book on evolution^14^. Since then, a multitude of descriptions of chondrodysplasias have been reported in sheep^15,16,17,18,19^. One such syndrome, characterised by either total or partial absence of distal phalanges in the limbs, achondroplasia of the head with protruding eyes, short ears and short tail, was first reported in South Country Cheviot sheep flocks in 1971^16^. In 2009, the presence of a similar phenotype, colloquially termed ‘stumpy’ lambs in North Country Cheviot sheep was brought to our attention by a sheep breeder (Supplementary Figure 1). The phenotype appeared identical to the previous report in South Country Cheviot sheep - the short tail and missing distal phalanges being particularly distinguishing factors. These lambs are generally culled shortly after birth, although there were also unconfirmed reports of deformed stillbirths. As reported earlier^16^, the condition was observed in offspring from specific rams and ewes that were phenotypically normal and in addition some cases were co-twinned to normal lambs. These observations suggested that the condition was under genetic control with a recessive mode of inheritance. Assuming that the affected individuals were identical-by-descent (IBD) for the causal mutation and shared a region of extended homozygosity around the causal locus, we scanned the genome for regions of homozygosity. Sequence analysis of the candidate region, under the assumption that affected lambs were homozygous and their putative carrier parents were heterozygous, identified a deletion leading to a premature stop codon in the *PRICKLE1* gene. We report the identification of this mutation and the characterisation of the associated phenotypes including multi-organ developmental defects similar to that observed in other PCP pathway mutants such as *Prickle1* mutant alleles in mouse^20,21,22^.

## Results

### Causative genetic variant maps to chromosome 3 - Figure 1

Farmer observations of a similar achondroplastic phenotype to that reported by Wray et *al*. (1971)^16^ (Supplementary Figure 1) occurring only in progeny from specific ram and ewe matings suggested that the condition could be genetic in nature. The parents of affected lambs were phenotypically normal. Furthermore, some cases were co-twinned to normal lambs, suggesting strong evidence for a genetic rather than environmental mechanism and indicative of a recessive genetic condition. DNA was isolated from tissue samples from euthanised affected lambs and appropriate controls and genotyped for single nucleotide polymorphisms (SNPs) using both the Illumina OvineSNP50 BeadChip and Ovine 800 HD BeadChip arrays.

**Figure 1.**
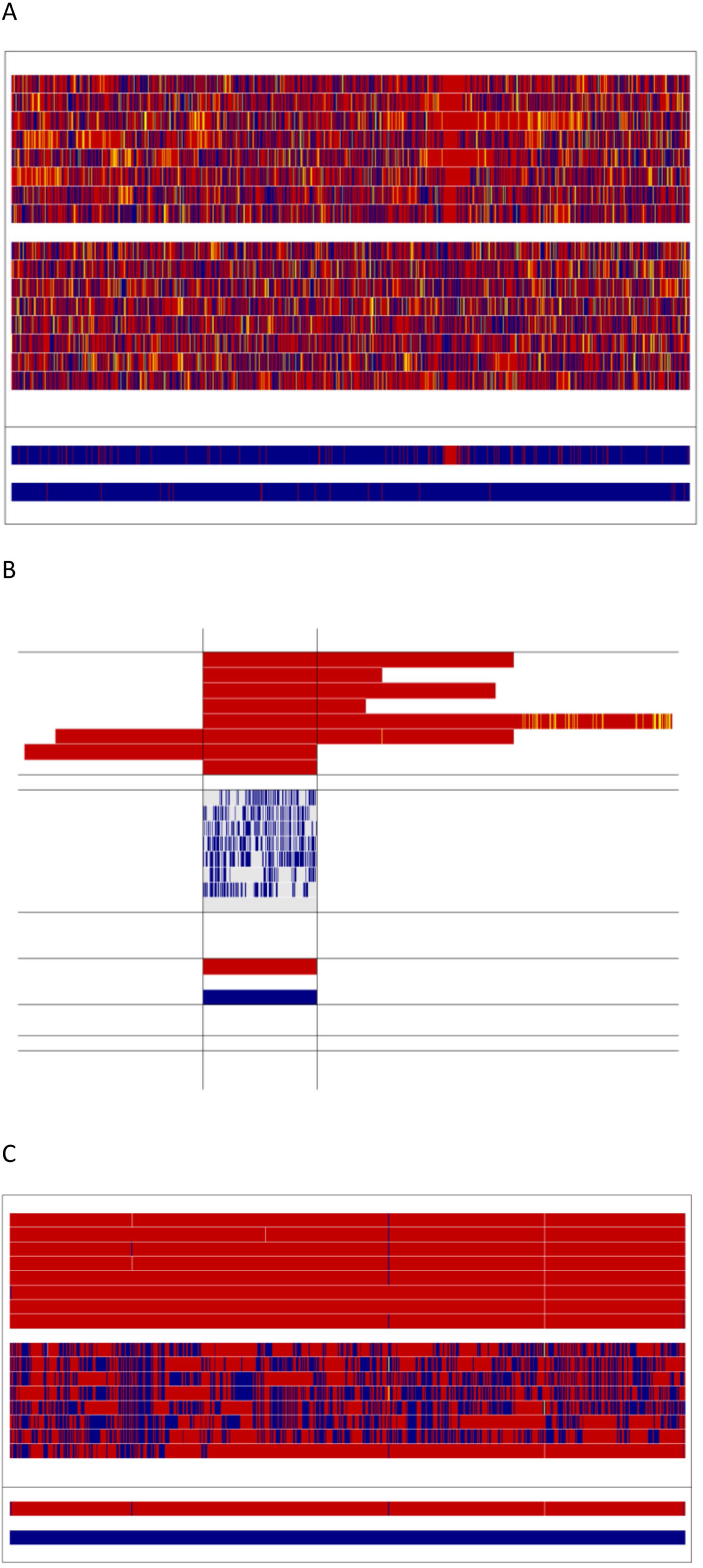
Homozygosity mapping. **A.** Homozygosity mapping of achondroplasia in Cheviot sheep assuming a single underlying recessive mutation – results for the full length of chromosome 3 (OAR3). Vertical blocks in red and yellow represent homozygous genotypes, and blue heterozygote genotypes. The eight cases (above) and eight parent controls (below) are shown (one per line). A summary of homozygosity mapping is provided cases vs control at the bottom of Fig 1A in which if all the animals within a group (*i.e.*, case or control) are homozygous for the same allele, then the relevant colour (red or yellow) is shown alternatively heterozygote SNPs are coloured blue. **B.** Extract from Fig 1A showing the longest ROH block in the cases on OAR3 ranging from rs419318162 to rs405796644, corresponding to position 142870240 – 147058642 in sheep genome sequence assembly Oar_v3.1 (https://www.ncbi.nlm.nih.gov/assembly/GCF_000298735.1/. The first eight lines are the cases; the second eight lines are the controls with genotypes shown only for the region of homozygosity shared across the cases. Genotypes shown for controls are shown only if they are different to cases (blank genotype in controls within the targeted segment means that they share the same genotype as cases). Finally, the bottom two lines are summary lines shown on the same basis as the summary lines in Fig 1A. **C.** A further magnification of the region of interest with genotypes coloured as in Fig 1A.

Mapping of extensive runs of homozygosity (ROH) in affected lambs in comparison with genotypes from independent unaffected animals as controls using the low-density SNP chip (Illumina OvineSNP50 BeadChip) and those from the high-density SNP chip (Ovine 800 HD BeadChip) using parents (putative carriers) as controls, identified a similar unique region on Chromosome 3 (OAR3) as being associated with this condition. The results from the high-density scan identified a broad ROH 4.2 Mb region with 734 SNPs, but with 2 discordant SNPs in the cases (Figure 1 A, B and C). Within the 4.2 Mb region, there were 3 ROH blocks. The first ROH block was from SNP rs419318162, (Oar_v3.1, 3: 142870240 bp) to SNP rs398321221 (Oar_v3.1, 3: 143640069 bp), inclusive, yielding a candidate region spanning 0.77 Mbp. The second ROH block spanned a 1.6 Mbp region from rs402985366 (Oar_v3.1, 3: 143651872 bp) to SNP rs417182779 (Oar_v3.1, 3: 145252109 bp), inclusive. The last ROH block of 1.8 Mbp was from SNP rs420216898 (Oar_v3.1, 3: 145275422 bp) to SNP rs411722153 (Oar_v3.1, 3: 147053343 bp) inclusive. [Note: all genome coordinates shown in parentheses above refer to sheep genome sequence assembly Oar_v3.1 (GCF_000298735.1]. One control animal shared the same genotype as the cases in the third block. The regions identified by homozygosity mapping above included homozygous SNPs at each end. Thus, the region to be subjected to fine mapping by sequence analysis was defined by the flanking non-homozygous SNPs, *i.e*., from SNP rs425896739 (Oar_v3.1, 3: 142862353 bp) to SNP rs405796644 (Oar_v3.1, 3: 147058642 bp).

### Sequence analysis reveals 10 bp deletion in *PRICKLE1* as candidate causative variant

To identify candidate mutations for the causative variant, whole genome sequence data from a pool of 5 cases and separately for 5 putative carrier dams were analysed (Supplementary Table 1). All variants called from the 5 putative carriers were merged giving a total of 30,690 SNPs/Indels (Indels are insertion/deletion variants) in the segment of interest on OAR3. The homozygous SNPs were 16,611 out of a total 16,992 raw variants in the pooled samples. However, only 727 of these homozygous pooled variants were heterozygous in the carriers. Analysis of the predicted SNP effects^23^ indicated the significance of a 10 bp deletion in the *PRICKLE1* gene (Figure 2). Other missense mutations were observed in the *ADAMTS20* gene. The rules were then relaxed to allow at least one heterozygous or missing (such as due to low sequencing coverage) variants when comparing the homozygous pooled variants to all the carriers. In this subset, we identified 15,956 common variants, other than *PRICKLE1*. The effects of all the other 12 missense SNPs were predicted to be low.

**Figure 2.**
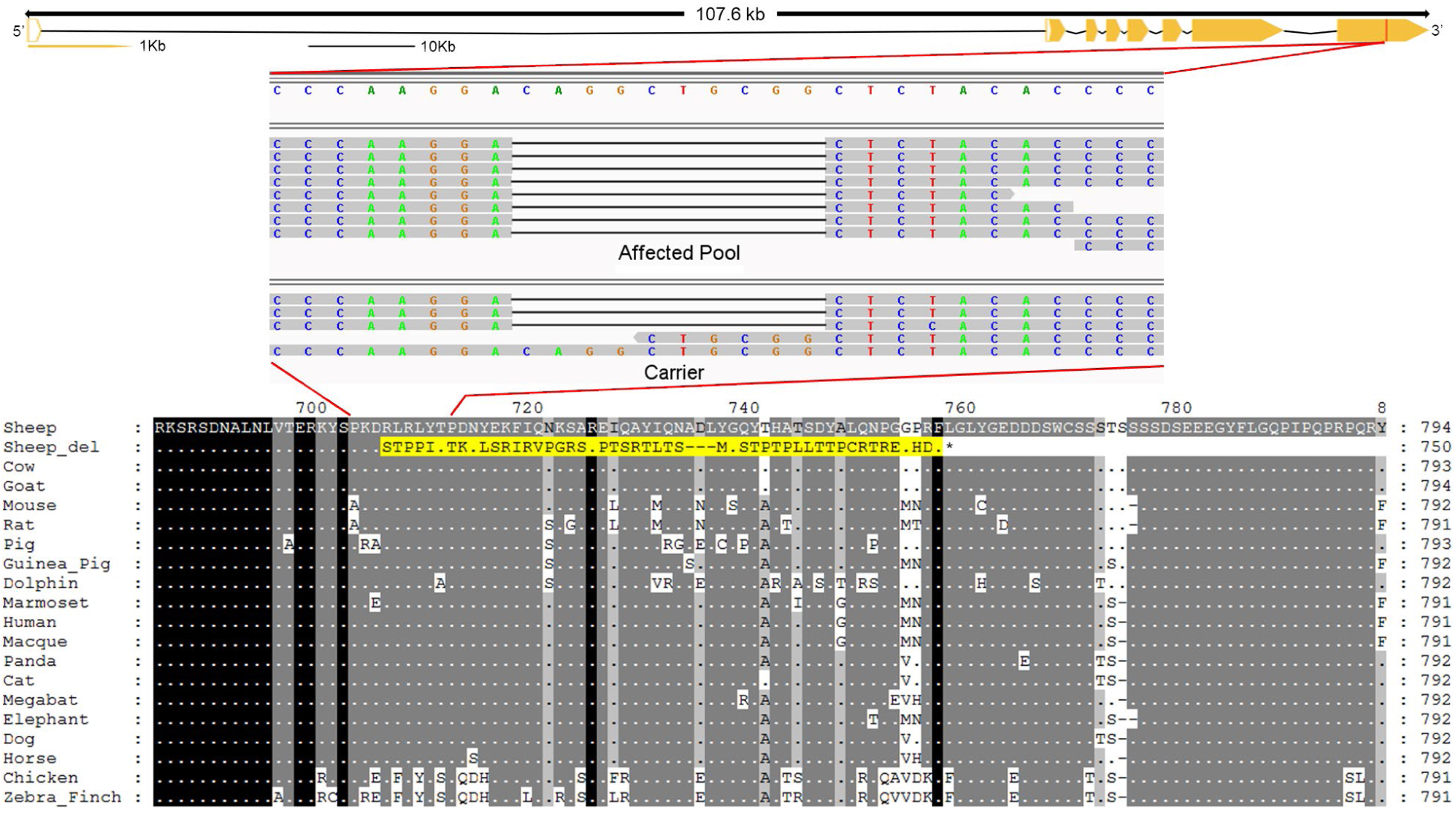
Identification of an indel mutation in the sheep *PRICKLE1* coding sequence. **A.** Characterization of *PRICKLE1* gene showing the position of the 10 bp putative causal deletion. Yellow block arrows indicate exons, filled arrows indicate coding sequence. Introns are indicated by a line. The position of the indel mutation in the gene is shown by a red vertical line. **B.** Expanded view of the region carrying the indel, with Integrated Genome viewer (IGV) tracks displaying reads from this region in a pool of homozygous affected lambs (upper) and from a single carrier (lower). The reference genome sequence is shown above the aligned reads. **C.** Multispecies translated sequence alignment showing conservation of amino acid sequence in this region of PRICKLE1.

To confirm that the 10 bp deletion in the *PRICKLE1* gene altered *PRICKLE1* transcripts, cDNA was prepared from tissues isolated from cases and carriers of the *PRICKLE1* deletion. RT-PCR fragments spanning the deletion site were subjected to Sanger sequencing (Supplementary Figure 2), that indicated that both wild type (WT) and mutant alleles are transcribed (Supplementary Figure 2). The 10 bp deletion and resulting frameshift affects the C-terminal region of PRICKLE1, replacing amino acids from position 702 to the terminus at 834 with 50 amino acids sharing no sequence similarity with any known protein.

At this stage in the study, a strong candidate causal variant (a 10 bp deletion in the coding sequence of *PRICKLE1*) had been identified based on homozygosity mapping and sequence analysis and similarity of the sheep phenotypes to those caused by *Prickle1* mutations in mice^20,21^. Phenotypic information on affected sheep was limited to farmer descriptions and comparisons with the phenotype reported previously^16^. Biological material was limited to ear tissue from euthanised affected lambs and DNA from putative carriers and unaffected animals in independent Cheviot populations with no history of the achondroplastic phenotype. Thus, additional affected lambs and appropriate controls were acquired for more detailed phenotypic characterisation.

### Gross anatomical phenotypes

A gross post-mortem examination by a veterinary pathologist was undertaken of lambs homozygous for the *PRICKLE1* deletion (n=9), and heterozygous carriers of the same deletion (n=3). All the heterozygous lambs were grossly normal. The gross anatomy of affected Cheviot lambs was very similar to the anatomy of the achondroplastic syndrome that was described previously^16^, including head, ear, eye, leg and tail malformations. All affected lambs (9/9) exhibited an abnormal stance, often resting on the forelimb carpus joint instead of standing (Supplementary Figure 1B), while parents and unaffected siblings stood normally. In all lambs, the coxofemoral, femorotibial and tibiotarsal joints were rigid and associated with marked, focally extensive articular cartilage loss at the lateral aspect of the tibial epicondyle and metatarsal condyle, as well as around the base of the ligament of the femoral head. This malformation resulted in an approximately perpendicular angle between the femurs and the pelvis. All affected lambs had bilateral hoof malformations on both forelimbs and hindlimbs, shorter snouts and tails that were short and stumped externally (Figure 3). In terms of forelimbs, three out of nine (3/9) presented two small and irregular hoofs, 2/9 a single irregular hoof, or only a pad of thickened skin 4/9. Hindlimbs were similarly affected; 5/9 presented two irregular hoofs, with a large medial hoof and a small lateral hoof, 4/9 two small and irregular hoofs. Snouts of affected lambs were also shorter than control siblings presenting a relatively shorter mandible (brachygnatia inferior -3/9) or a relatively longer mandible (prognatia inferior -1/9). In addition, 3/7 affected lambs had some degree of hydronephrosis characterized by variable dilatation of the renal pelvis, and mild hydronephrosis was seen in two lambs, whereas the other lamb had moderate hydronephros and unilateral hydrourether. Five out of nine (5/9) lambs had hair on the forehead that was curly and convoluted, a pattern that was not observed in any of the control lambs. One out of the nine affected lambs (1/9) also exhibited spina bifida, having a segmental loss of the spinous processes of all the lumbar vertebrae, associated with lack of paravertebral musculature on the dorsal aspect of this area, extending caudally over the coccyx (spina bifida oculta). One out of the nine affected lambs (1/9) had protruding eyeballs (exophthalmia), which was associated with bilateral corneal ulceration and anterior chamber prolapse. The frequencies of the defects observed at post-mortems of affected lambs are summarised in Supplementary Figure 3.

**Figure 3:**
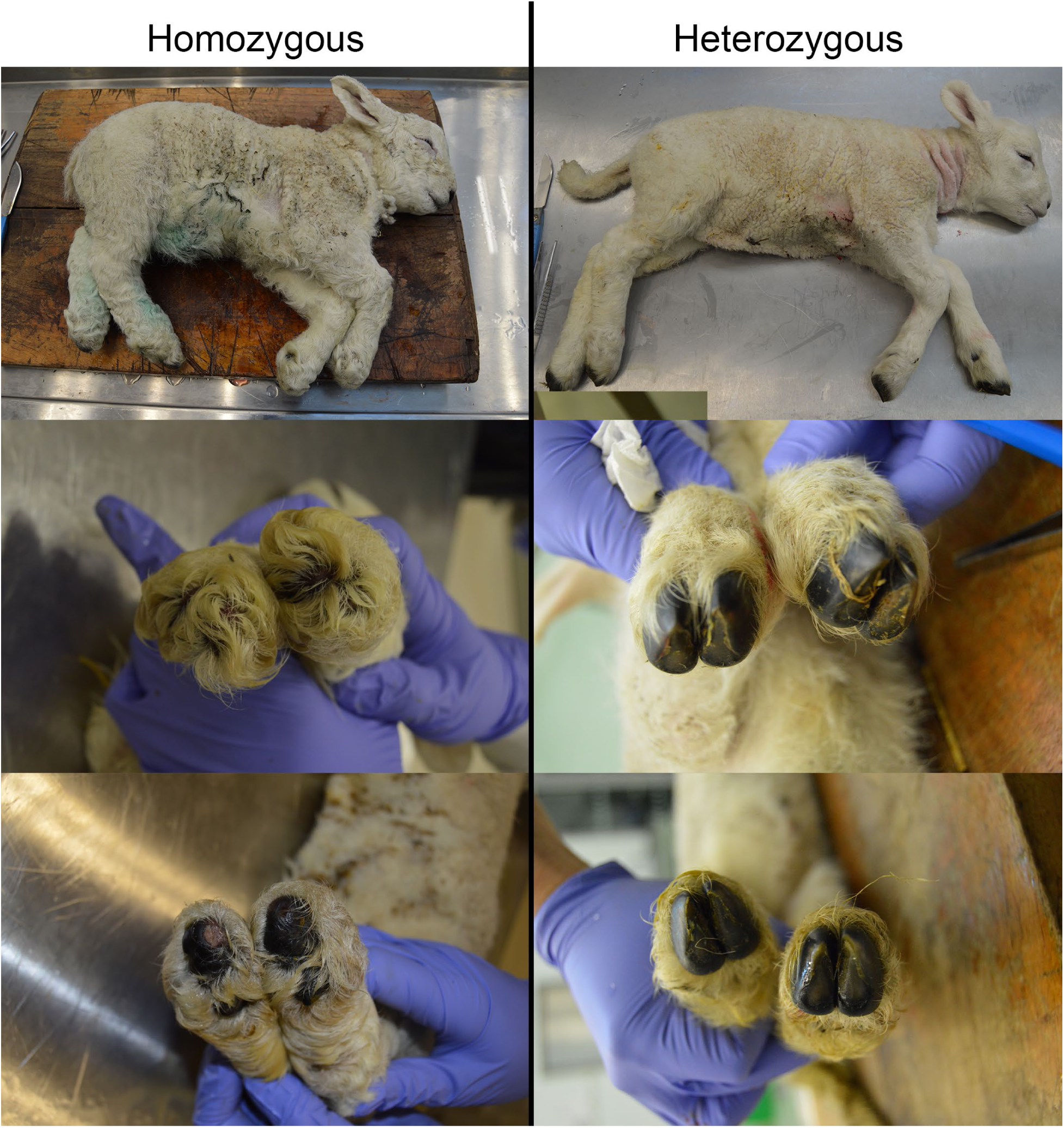
Gross phenotypes. Gross presentation representative of *PRICKLE1* 10 bp deletion in homozygous vs. heterozygous lambs. The rows feature: (top) a lateral view of the whole body, (middle) forelimb hoofs, and (bottom) hind limb hoofs. The gross presentation of heterozygous lamb (right column) is within normal limits, whereas the homozygous lamb (left column) presents with several developmental abnormalities. These abnormalities include disproportionate dwarfism, dome-shaped calvaria, relatively long pinna and short snouts, short and stumped tail, and bilateral hoof malformations of all limbs.

### Skeletal assessment by Computed Tomography (CT)

Computed Tomography (CT) analyses of eight affected and three age-matched control lambs was undertaken [Supplementary Table 2: lambs CH1, CH2, CH3, CH6, CH10-17]. CT analyses revealed no skeletal abnormalities in the three age-matched control lambs (CH3, CH10, CH11) and vertebral column abnormalities in the nine affected lambs (CH1, CH2, CH6, CH12-CH17 inclusive), of which six had multiple abnormalities. These abnormalities included one T13 transitional vertebra, three lambs with deviation from the normal vertebral distribution of 13 thoracic and 6 lumbar vertebrae, three cases of sacral and/or coccygeal spina bifida, one L1 cleft vertebra, five lambs with hemivertebrae, in four of them multiple (2 x T3, 1 x T4, 2 x T5, 2 x T7, 1x L1) and presumed secondary perihemivertebral scoliosis (Figure 4).

**Figure 4:**
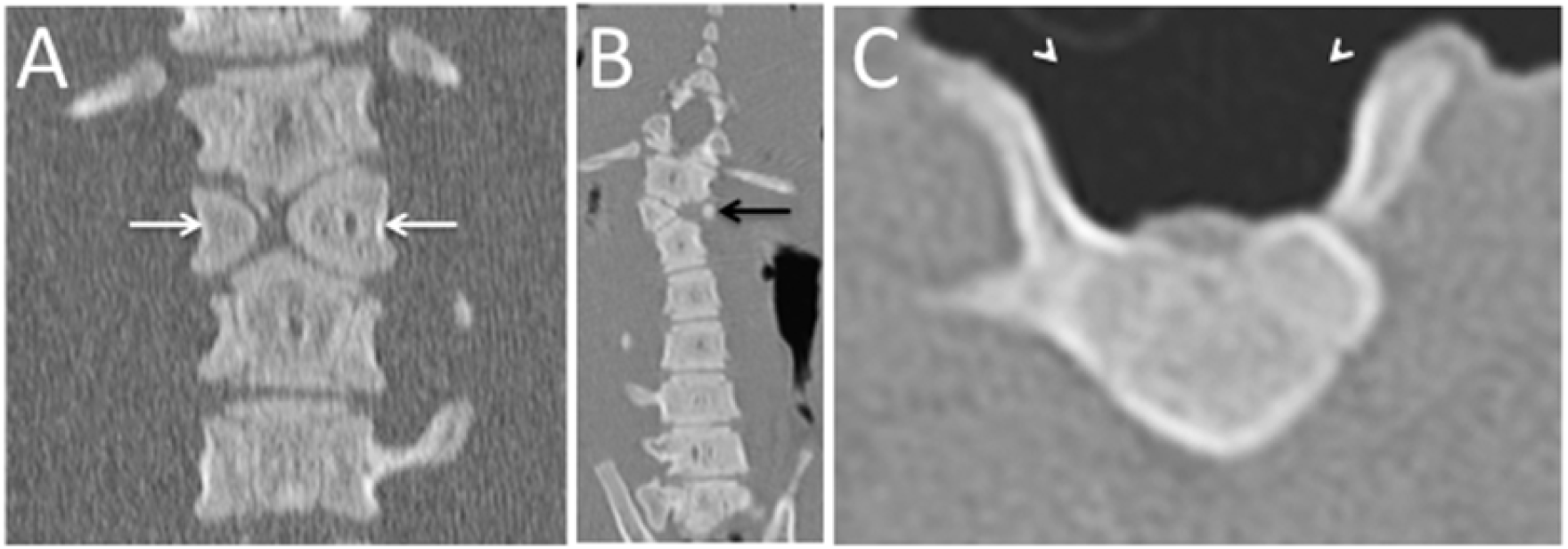
Computed Tomography images of vertebral malformations in *PRICKLE1* mutant lambs: A) Dorsal slice image of a L1 cleft vertebra resulting in two funnel shaped vertebral body halves (white arrows). B) Dorsal slice image of the lumbar vertebral column showing a missing left vertebral body half (black arrow), an opposing wedge-shaped right vertebral body half and secondary vertebral curvature deformation (scoliosis). C) Transverse slice image of a L2 spina bifida malformation in which the dorsal arch is missing (arrowheads).

### Craniofacial phenotypes

Three-dimensional reconstructions of the CT scans of ten skulls (seven mutants, three carriers) were used to assess face shape. The skulls of the affected lambs that were homozygous for the 10 bp deletion in *PRICKLE1* were large and dysmorphic. The facial shape of affected lambs differed markedly from carriers. Each of the mutants showed obvious signs of facial retrusion (Figure 5 A-D). Using a landmark-based geometric morphometrics analysis, we characterized the morphospace of the skulls, focusing on the facial bones. Nine landmarks were placed across or adjacent to the viscerocranium (Figure 5 A-D). Due to the limited sample size, we were unable to determine whether the scale of the face differed between Stumpy mutants and carrier siblings; log (centroids), the proxies for the amount of scale applied to the Procrustes fit, were nominally different when unequal variances were assumed (two-sample Student’s t-test: heteroscedasticity, *p*=0.05; homoscedasticity, *p*=0.14). Principal components analysis (PCA) was used to decompose the covariances of shape that remain following the Procrustes fit. To investigate whether allometry differs between Stumpy and carrier lambs, we compared the regression scores against the log (centroid size). Grossly, intercepts differed by genotype, suggesting that shape differences could be due to allometry (data not shown). Regardless of whether we included allometry or removed its effects on shape, two distinct clusters of facial shape data were observed with cluster membership corresponding to genotype (Figure 5 E-F). We conclude that facial retrusion occurs in affected lambs, a result that parallels finding in a mouse *Prickle1* mutant^21^. Our analysis suggests that the disparity between the facial skeletons of Stumpy mutants and carriers is due to alterations in morphogenesis that affect bone shape.

**Figure 5:**
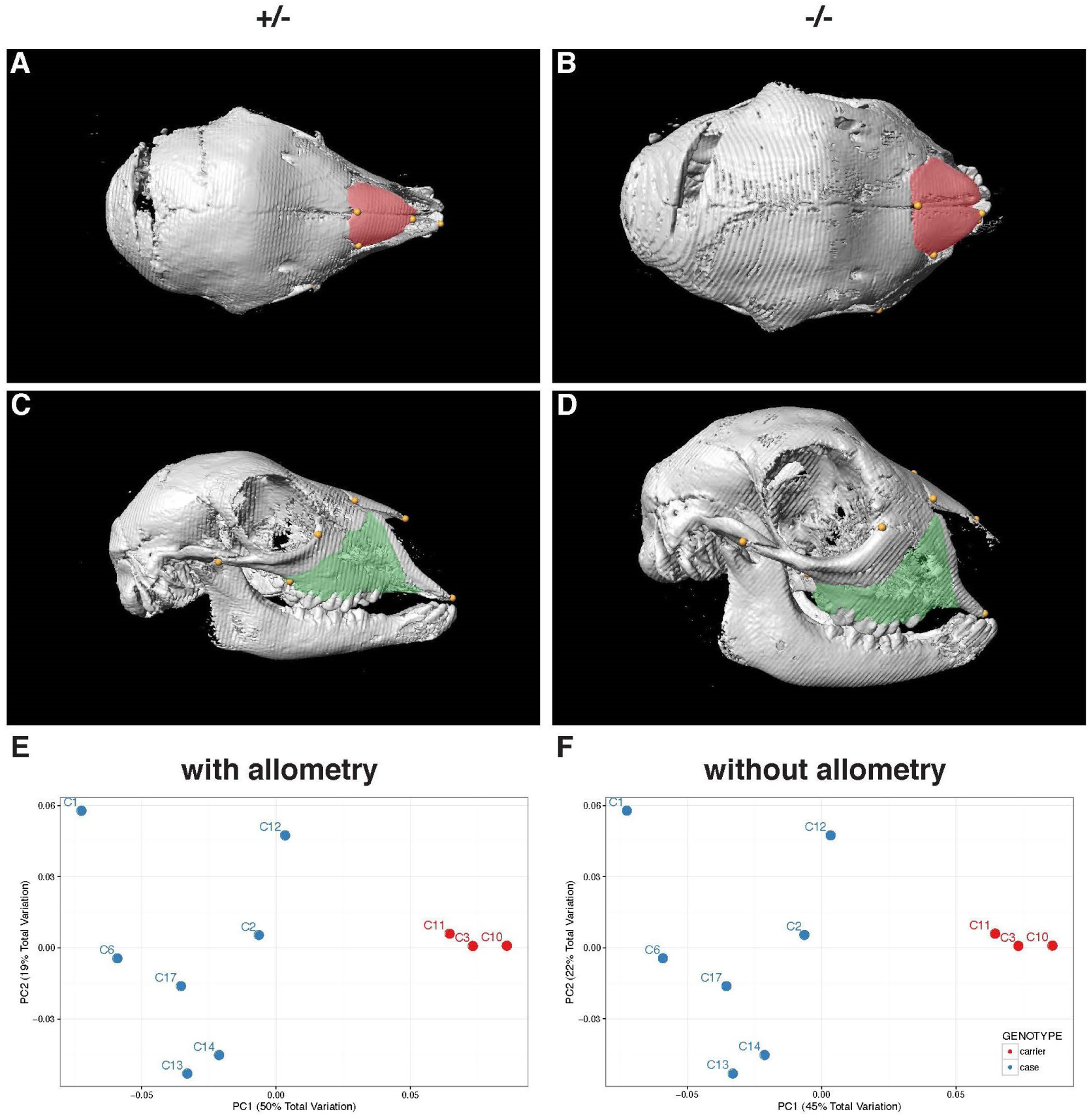
Morphometric analysis of skull reconstructions of *PRICKLE1* mutant lambs. Skulls reconstructed from CT scans (A-D). Representative dorsal (A, B) and lateral (C, D) views of isosurfaces from the affected lambs that are homozygous for the 10 bp deletion in *PRICKLE1* (B, D) and age matched carrier lambs (A, C). From the dorsal view, the nasal bone (pseudo-colored red) of mutants is broad and spade-like (compare B to A). In the lateral view, much of the mutant’s caudal maxilla (pseudo-colored green) is tucked under the orbits (D) compared to carriers (C). Note that damage to the cranial vault is an artifact of postmortem tissue sampling. Bivariate plots of PC1 and PC2, with (E) and without (F) allometry illustrates the clean separation of face shapes according to genotype.

### Bone growth morphology

Gross measurements of the tibiae and CT reconstruction of tibia, radii, and metatarsi demonstrated that the tibiae of affected lambs were both shorter (p = 0.017) and wider (p = 0.025) than those of carrier siblings (Supplementary Figure 4, Supplementary Table 3). Histological evaluation of H&E staining of multiple sections from sibling lambs (CH2, CH3; 49 and 57 sections respectively) revealed a significantly smaller total growth plate length (994.12 µm (se 19.32 µm) versus 1160.34 µm (se 17.39 µm), p<0.0001, Figure 6) and smaller proliferating zone (PZ; orange bar) in the Stumpy lamb (individual CH2) (256.46 µm (se 6.92 µm) versus 401.49 µm (se 6.22 µm), p<0.0001, Figure 6), although the resting zone (RZ) and hypertrophic zone (HZ) were comparable with the CH3, WT lamb (Figure 6) (Supplementary Table 4). Chondrocytes within the PZ appeared similar between the lambs, although there was some loss of organisation and polarity in CH2, consistent with a milder phenotype than has been observed in mice with a *Prickle1* KO phenotype, where the chondrocytes are highly disordered and the achrondroplasia is more severe (Figure 6). We conclude that the skeletal dysplasia characteristics exhibited by mutant lambs are highly similar to unique features of disrupted PCP signalling observed in *Prickle1* knock-out mice and other PCP mutants, supporting the conclusion that the 10 bp deletion in *PRICKLE1* is responsible for the skeletal dysplasia phenotype in affected lambs.

**Figure 6:**
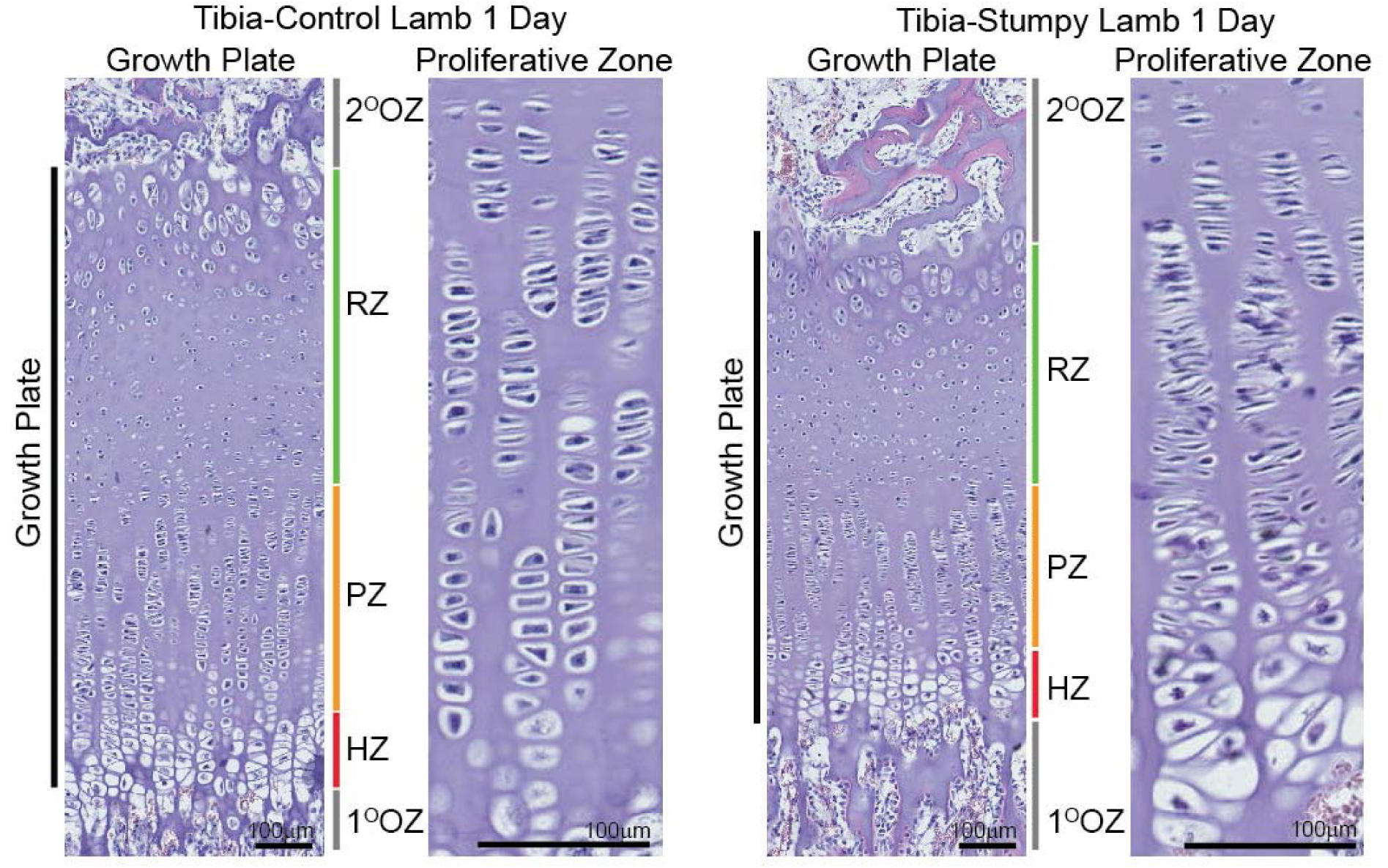
Bone growth morphology. Haematoxylin and eosin-stained sections of the tibia growth plates from 1-day-old case (stumpy) and control lambs. The chondrocytes situated in both the proliferating (PZ) and hypertrophic (HZ) zones of the growth plate were stacked in columns in the direction of growth and their organisation and polarity were similar in the control and case lambs. The higher magnification micrographs of the proliferation zones clearly show the arrangement of cells within the chondrocyte columns. 2^O^OZ = secondary ossification zone, RZ = resting zone, PZ = proliferative zone, HZ = hypertrophic zone, 1^O^OZ = primary ossification zone. Scale bars = 100 µm

### Localisation of wild-type and mutant PRICKLE1 protein

We found that the mutant allele produced a transcript (Supplementary figure 2) and assessed the presence and location of the mutant PRICKLE1 protein. An antibody raised against PRICKLE1 amino acids (aa) 321-580, of the ∼830aa in human PRICKLE1, was used for immunohistochemistry analyses of sections of trachea. In 1/1 control and 1/1 affected samples PRICKLE1 was observed in the respiratory epithelial cells of the trachea, indicating that the wild-type and mutant transcripts were both translated into protein. In the trachea cells from the affected lamb the PRICKLE protein was localised to the apical surface, whereas localisation of the protein was more diffuse in control cells (Figure 7 D and C, respectively).

**Figure 7:**
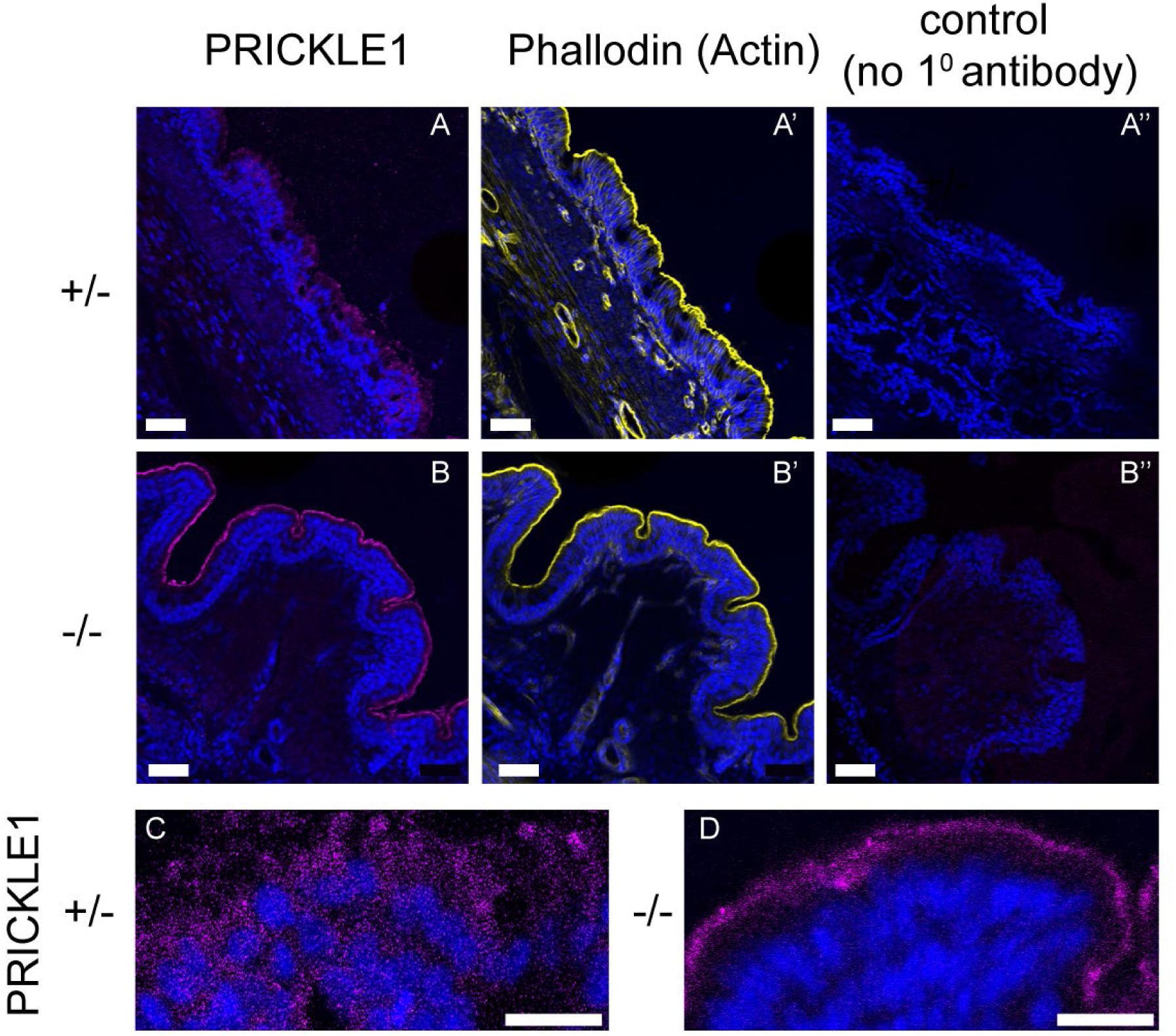
Localisation of PRICKLE1 wild-type and mutant protein in lamb tissue. A-D Anti-PRICKLE1 immunofluorescence detecting the protein in (A, C) heterozygous (+/-) control and (B, D) homozygous mutant (-/-) tracheal epithelium of 1-day old lambs (sibs). PRICKLE protein detection is indicated by the purple signal. Tissue structure and apical-basal polarity of respiratory epithelium are indicated by F-Actin detection (phalloidin, yellow) in A’ and B’. Negative control samples without primary antibody are shown in A” and B”. Nuclei are stained blue by DAPI in all panels. Scale bars (white): A-B”, 50 μm; C, D 20 μm.

Many aspects of the abnormal anatomy of *PRICKLE1^-/-^* lambs can be common to failures in several different genetic pathways or are known to be multi-factorial. In addition, other than short stature, presentations of the range of abnormalities were variable between individuals, and some, including spina bifida and corneal ulceration were rarely observed. In addition, to a reduction in long bone length, *PRICKLE1^-/-^* lambs’ legs were shortened by the failure of phalanx development exhibiting a variable loss of phalanges in both forelimb and hindlimbs (Figure 8 A-F). The bilateral loss of distal limb formation specifically effecting the development of the phalanges, a form of brachydactyly was present in all *PRICKLE1^-/-^* lambs; 8/8 *PRICKLE1^-/-^* showed bilateral loss of digits. 2/3 *PRICKLE^+/-^*lambs had normal phalanges (see Figure 8 for examples).

**Figure 8:**
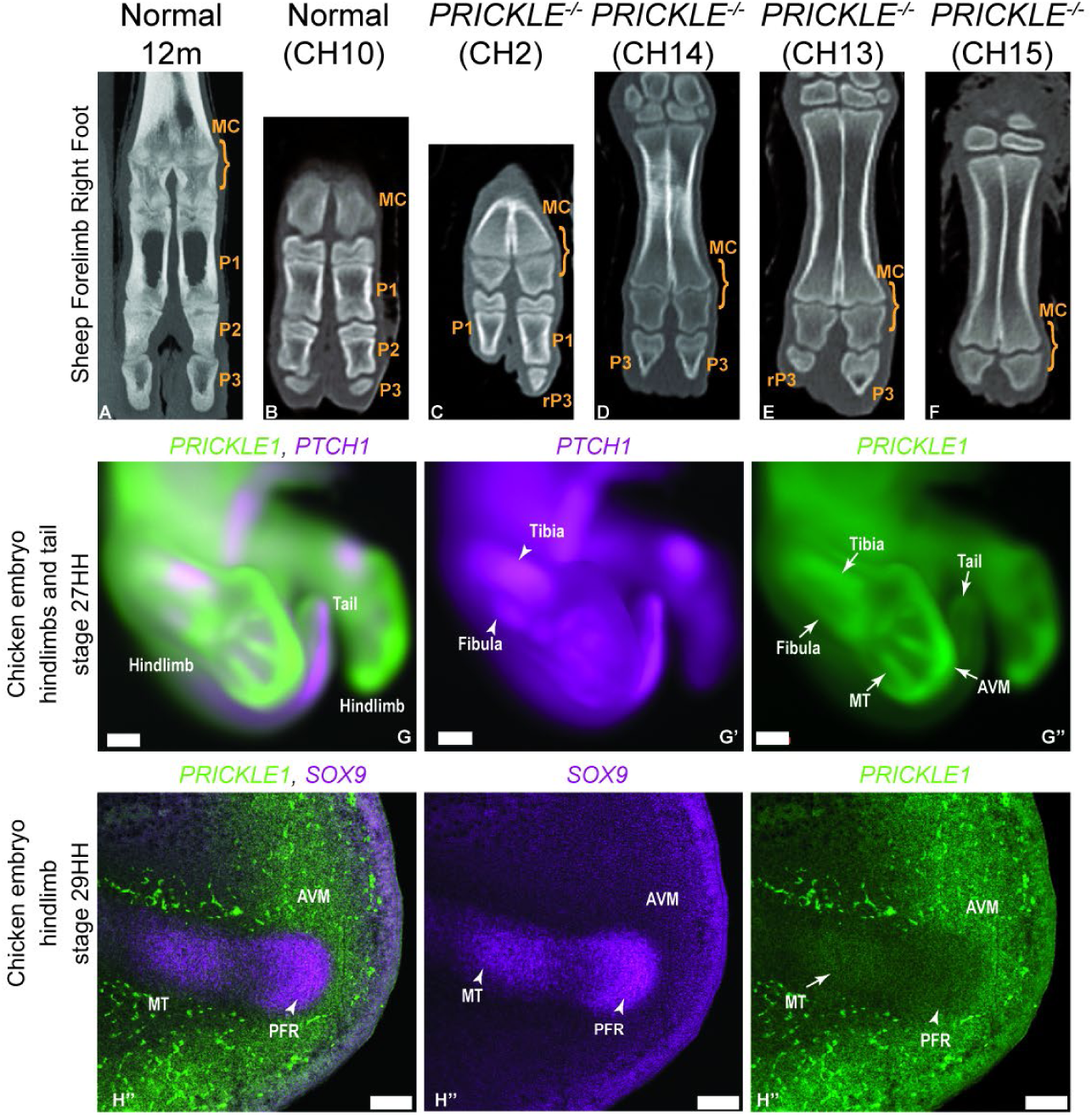
Lamb toes and *PRICKLE1* RNA expression in chicken embryos. A-F: ‘Stumpy’ PRICKLE1-/- lambs show loss of phalanx formation in distal leg. A. Forelimb of a normal sheep at 12 months of age, demonstrating digits 3 and 4, the normal digits of a sheep, consisting of a proximal fused metacarpal, and distally phalanx 1, 2, 3. B. An unaffected lamb (CH10), showing the same pattern proximal fused metacarpal, and distally phalanx 1, 2, 3. Affected lambs (C-F) showed intact metacarpals but a variable loss of phalanges, including (C) a loss of P2 and 3, (D, E) a loss of P1 and P2 or (F) a total loss of P1, P2 and P3. Defects could be different in toes of the same foot (C, E) and in bilateral limbs, and occurred in both forelimb and hindlimb. Right forelimb foot of all animals, shown as example. MC=metacarpal, P1=phalanx1, P2=phalanx2, P3=phalanx3, rP3- residual phalanx3 G-H”: *PRICKLE1* RNA expression localises to areas of altered anatomy in chicken embryos. *PRICKLE1* expression (green) is observed in chicken embryos in areas associated with developmental defects in *PRICKLE1*^-/-^ lambs, including the long bones such as the tibia and fibula and metatarsals (arrows, G’’, H’’), where it is co-expressed with *PTCH1* (magenta), in chondrocytes undergoing endochondral ossification and in cartilage areas in advance of ossification. *PRICKLE1* expression is also observed in the tip of the tail, an area associated with the growth zone of the tail (arrow G’’). In the developing digits, *PRICKLE1* expression is seen in *SOX9^+^* chondrocytes (magenta) undergoing endochondral ossification (arrow, H’’), but not in the *SOX9^+^* chondrocytes of the ‘phalanx forming region’ (PFR). In contrast, *PRICKLE1* expression is observed in the avascular mesenchyme (AVM; G’’, H’’), and is particularly enriched over the PFR (G’’), a tissue that will contribute to the newly forming phalanges and that is subject to high WNT5a signalling. Scale bar G= 500 µm, H=100 µm MT= metatarsal, PFR= phalanx forming region, AVM= avascular mesenchyme

In order to examine if the loss of digits is associated with the loss of long bone development or an independent mechanism relating to digit development, we undertook Hybridisation Chain Reaction (HCR) RNA *in situ* hybridisation in chicken embryos, that are commonly used as a model of vertebrate limb development. *PRICKLE1* expression was observed in association with *PTCH1* in the tibia and fibula long bones during endochondral ossification of the cartilage primordia, and in advance of endochondral ossification, in the metatarsal cartilage primordia, prior to *PTCH1* expression (Figure 8 G; arrows indicate *PRICKLE1* expression). In addition, we observed *PRICKE1* expression at the tip of the tail and throughout the distal limb mesenchyme specifically enriched distally to the digital ray. To examine this further, we examined expression of *PRICKLE1* in association with *SOX9*, that is expressed in cartilage primordia and in the ‘phalanx forming region’ (PFR), an organising tissue essential for phalanx development and patterning (Figure 8 G – H”). *PRICKLE1* was expressed in the developing metatarsal phalanx in association with *SOX9*, but not in the PFR. Instead *PRICKLE1* expression was observed in the ‘avascular mesenchyme’ (AVM), the population of cells that contain the progenitors of the rest of the digit.

## Discussion

Inherited disease represents a burden for the sheep and wider farmed animal sector, but the associated naturally occurring mutations can offer opportunities to study gene functions in animals other than laboratory rodents. Genetic diseases with a recessive mode of inheritance can be a particular challenge for the animal breeding sector, especially in dairy cattle where large geographically dispersed sire families can be established before the disease is detected and a carrier male identified^24^. A growing catalogue of inherited diseases in which the causative mutation has been identified is available in the Online Mendelian Inheritance in Animals (OMIA)^25,26^, to which a record of the *PRICKLE1* mutation described here has been added (OMIA:002693-9940). DNA diagnostic tests are available for many of these diseases to facilitate the eradication of these disease burdens.

A skeletal abnormality in North Country Cheviot lambs was brought to our attention by a breeder. Affected lambs exhibited rounded heads, short legs, a loss of hooves, abnormal stance and short twisted tails and were referred to as ‘stumpy lambs’. As the condition had been observed in lambs from specific ram and ewe matings with cases observed co-twinned to normal siblings, it was likely that the syndrome was caused by a recessive genetic condition. Individuals affected by rare recessive mutations are likely to share chromosomal regions around the causative genetic variant that are identical-by-descent and exhibit extended runs of homozygosity. Thus, homozygosity mapping even in small numbers of affected individuals is a powerful method for mapping such recessive mutations^27,28,29^. We used homozygosity mapping initially in affected lambs and controls from an unrelated population and subsequently in affected lambs and putative carrier parents to map the causative variant to a 4.2 Mbp region on chromosome 3 (OAR3) (Figure 1). Through subsequent genome sequence analysis of this region on OAR3 we identified a 10 bp deletion causing a frameshift in the open reading frame of *PRICKLE1* a gene essential for normal vertebrate embryonic development (Figure 2).

The identification of this compelling causative mutation enabled the development of a simple DNA diagnostic test that was made commercially available and facilitated the removal of carrier rams from the breeding population.

Having identified the 10 bp deletion in *PRICKLE1* as a strong candidate causative variant we acquired additional material from affected lambs and appropriate controls in order to i) test the association between *PRICKLE1* genotypes and the stumpy phenotypes and ii) more fully characterise the phenotypes at multiple levels. A further 8 affected stumpy lambs were found to be homozygous for the 10 bp deletion in *PRICKLE1* and 32 parents of affected lambs found to be heterozygous for the 10 bp deletion in *PRICKLE1*, *i.e.*, proved to be carriers of the mutation.

The presence of both wild type and mutant transcripts was detected at an approximately equal ratio. The frameshift caused by the deletion is in the last exon of the gene, and so would be predicted to spare the transcript from nonsense mediated decay (NMD)^30^, permitting translation of the mutant protein. Indeed, we detected PRICKLE1 protein in tracheal epithelial cells of an affected lamb that was homozygous for the 10 bp deletion in *PRICKLE1* (Figure 7). However, the mutant PRICKLE1 protein appeared to be mis-localised in those cells. The mutant protein is predicted to be missing the entire C-terminal PRICKLE1 homologous domain (PKH) that uniquely identifies PRICKLE family proteins, while sparing the N-terminal PET domain and three LIM domains that are found in other related proteins^31^. The N-terminal PET and LIM domains mediate specific protein-protein interactions^32^, while the C-terminal region absent from the mutant protein carries a D-box motif involved in protein degradation and modulation of cell signalling^33^ and a farnesylation site involved in association of PRICKLE1 with cell membranes^34^ and regulation of cell migration^35^.

It is likely that the protein encoded by this mutant sheep allele retains some function. Early embryonic lethality has been reported for mice with a complete null allele of *Prickle1*^36^. However, Liu *et al*. (2014)^21^ reported survival to postnatal days 2-4 in mice with *Prickle1* null mutations and attribute the difference in survival to the genetic background on which the mutations were created. A truncation mutation in mouse *Prickle1* that spares the PET and two LIM domains permits survival to birth, with short limbs and face, and deformed vertebrae^20^.

The results from the morphometric analysis are consistent with the idea that PRICKLE1 is required for cell polarization, particularly as it is needed for facial outgrowth. While there was little evidence that the facial bones between mutants and carriers were of different size, the shape space occupied by the facial bones of mutants is distinct from that of heterozygous carriers (Figure 5). Our observations are suggestive that in Stumpy mutants enough tissue was specified for facial development, though presumably defects in polarization resulted in arrest of the proximodistal extension of the face.

The results from the HCR RNA expression analyses in developing chicken embryos (Figure 8 G-H) suggest that *PRICKLE1* has at least two functions in limb development, one early in the process of endochondral ossification of the cartilage primordia to bone, and one in the formation and patterning of the phalanges, independent of functions in other long bones. We have confirmed this in subsequent work^37^.

Phenotypes associated with a loss of *PRICKLE1* function have been described in both humans as well as model organisms including mouse, *Xenopus* and zebrafish models (reviewed^38^) and include spina bifida, lumbosacral myelomeningocele, microcephaly, hydrocephalus, heart and kidney defects, kyphoscoliosis, fused and missing ribs, under developed genitalia, body elongation defects, craniofacial defects including cleft palate and brachydactyly. Bilateral loss of phalanges that is a form of brachydactyly is rarely observed in humans or animals, but is observed in the null allele of the *Prickle1^-/-^* mouse^20^ and is a defining phenotype of human Robinow Syndrome (MIM #180700, *WNT5a*; MIM #268310, *ROR2*), particularly that caused by recessive mutations in *ROR2*^39,40^. Robinow syndrome is a complex human genetic condition, that is associated, not with *PRICKLE1* directly but with other associated genes in the PCP pathway. Currently, while GWAS studies have suggested that *PRICKLE1* is associated with neural tube defects^41^ and craniofacial development^42^ there are, as yet, no reported human patients with loss of function mutations in *PRICKLE1* who have these phenotypes.

## Materials and Methods

### Animals

In 2009, the original animal material comprised ears harvested post-mortem from 8 Cheviot lambs that exhibited signs of achondroplasia. The controls were obtained by using 27 Cheviot Thoka project population available from independent flocks and previous study^43^. During the 2015 lambing season additional samples comprised affected lambs, age-matched unaffected lambs and nasal swabs from parental individuals from four farms (Supplementary Table 2). A further 162 sheep and 33 rams were nasal swabbed in flocks that had reported incidences of affected lambs. Hy-Line chicken lines were maintained under Home Office License PP9565661 at the National Avian Research Facility at the Roslin Institute, University of Edinburgh.

### Genotyping

DNA isolated from affected lambs and controls were genotyped for single nucleotide polymorphisms using the Illumina OvineSNP50 BeadChip array (https://emea.illumina.com/products/by-type/microarray-kits/ovine-snp50.html) and the Illumina Ovine 800 HD BeadChip^44^.

### Homozygosity mapping

Genotypes were obtained from 8 cases (putative homozygotes for the causative allele) and 27 unaffected sheep from an independent population from the same breed (putative heterozygotes or homozygous wild-types) using the Illumina OvineSNP50 BeadChip array. As validation, the Illumina Ovine 800 HD BeadChip was used in homozygosity mapping on genotypes from 8 cases (putative homozygotes for the causative allele) 1 sire and 7 carrier dams (putative heterozygotes). Only those SNPs (out of 606,006) that mapped to known positions on autosomal chromosomes and were not fixed nor completely heterozygous were retained. This filtering step left 52,0274 segregating autosomal SNPs for the use in homozygosity mapping with the HD chip. SNP positions within the Oar_v3.1 (GCA_000298735.1) genome sequence were used for the analyses.

### Sequencing

DNA from a pool of 5 cases and individual DNA samples from 5 dams were sequenced. Edinburgh Genomics used their standard GATK pipeline to call variants in each sample. For the pooled sample of 5 sheep the UnifiedGenotyper tool was used setting ploidy to 10 and for the other samples the HaplotypeCaller was used. Reads were aligned to sheep reference genome sequence version 3.1 (Oar_v3.1, GCA_000298735.1). Variant filtering required candidate variants to fall within a region of interest (OAR3 142862353 to 147058642) and met both the following criteria:

A. Completely homozygous for the non-reference allele in the pooled sample of cases
B. Heterozygous in each individual sample for which there is sufficient coverage to have confidence in the genotype. If the variant was homozygous in individual sample(s) that had low coverage at that position, the variants were not be ruled out since there was still the possibility of them being heterozygous.

### Mapping and variant calling

Raw data were aligned to the sheep genome sequence assembly Oar_v3.1 (https://www.ncbi.nlm.nih.gov/assembly/GCF_000298735.1/) using BWA-MEM 0.7.10 with default parameters^45^. Duplicates were marked using Picard tools version 1.115 (https://sourceforge.net/projects/picard/files/picard-tools/1.115/). ReadGroup information was added to the files using Picard tools version 1.115.

Variants were called using the GATK analysis pipeline in accordance with their recommended best practices. Further details can be found at http://www.broadinstitute.org/gatk/guide/ :

1. Local realignments of the BAM files were performed using GATK analysis toolkit version 3.2-2. Duplicates were marked with picard-tools version 1.115
2. Base quality recalibration was performed using GATK analysis toolkit version 3.2-2 using dbSNP sheep SNPs (downloaded 28/09/14) as a source of known SNPs
3. Variants were called on individual samples using GATK analysis toolkit version 3.2-2 HaplotypeCaller and UnifiedGenotyper
4. Variants were called across all samples simultaneously using the GATK analysis toolkit version 3.2-2 GenotypeGVCFs tool
5. Variant recalibration was performed using dbSNP sheep SNPs (downloaded 28/09/14) and sheep SNPs from Ensembl (downloaded 04/11/14).
6. Effects of identified variants were predicted based on the annotation of Oar_v3.1, ENSEMBL release 76, using the tool SnpEff v4.0 ^23^

### Phenotypic and post-mortem characterisation of cases

To further understand the disease, descriptive characterisation of the affected animal together with post-mortems were conducted. Gross post-mortem examinations by a veterinary pathologist were undertaken of lambs homozygous for the PRICKLE1 deletion (n=9), and heterozygous carriers of the same deletion (n=3). Internal organs were unavailable for examination in two of the heterozygous lambs (due to predation). Measures were taken of the head length (*i.e.*, from base of head to tip of philtrum), and pinna length (*i.e.*, from the base to the tip). The pinna length/head length was then compared between groups. The data were tested for normality using an Anderson/Darling normality test with means of measurements between homozygotes vs. heterozygotes compared using a two-sample t-test. The threshold for significance for this test was set at p=0.05.

### Computed Tomography (CT)

Animals were CT scanned to provide an accurate description of bone and cartilage morphology and shapes in cases vs control. Computed Tomography (CT) was performed on lambs post-mortem using a helical 4-slice CT unit (Somatom Volume Zoom, Siemens, Germany), including a whole-body series with 3 mm slice width and 1 or 2 mm slice width series of the head and selected long bones. The CT settings used included: 120 kV tube voltage, 76 to 208 mAs adaptive tube current, 1 or 1.5 s tube rotation time, 207 to 380 mm display field of view and 512^2^ image matrix. Images were reconstructed into DICOM format with bone and soft tissue kernels (proprietary names H40, B40, H70, B70).

All CT studies were retrospectively reviewed for skeletal abnormalities by a board-certified veterinary radiologist (TS), who was blind to the disease status of the animals at the time of evaluation. Images were evaluated using dedicated DICOM viewer software (Osirix, Geneva, Switzerland, version 5.8.5-64bit) on a computer workstation (Mac Pro, Apple, USA) with a calibrated LCD flat screen monitor (Thunderbolt Display, 27 inch, Apple, USA). During the course of image evaluation, multi-planar reconstructions and variable windowing were used according to the preferences of the viewer. Standard anatomic and diagnostic imaging reference were used for the evaluation^46,47,48,49^.

A multivariate analysis of landmarks was conducted on CT scans of carrier and affected sheep. Briefly, DICOMs were imported into Stratovan Checkpoint (v2015.06.30.0350 WIN x64) and each skull was converted to a surface file (*.ply). Nine landmarks were placed on each skull surface at locations across the viscerocranium. Using a custom R script, the Cartesian coordinates were captured and loaded into MorphoJ (v1.06a). A full Procrustes fit was used to translate, rotate, and scale the bulk data. A covariance matrix was generated from the Procrustes coordinates. To reduce the dimensionality of the data, a principal components analysis was run on the covariance matrix, providing an eigendecomposition that describes the shape variation in dimensions of successively smaller explanatory size. Allometry was removed by regressing the Procrustes coordinates over the log (centroid). A covariance matrix was generated from the residuals and analyzed by PCA. For illustrative purposes, skull reconstructions from two female sheep (postnatal day 3) were exported from CheckPoint and pseudo-colored using Photoshop (v13.0.6 x64).

### Bone growth morphology

Proximal tibia were dissected from 9 lambs and fixed in 10% neutral buffered formalin (NBF) for one week. Further processing was undertaken blind of genotype. Bones were decalcified in 10% EDTA, pH 7.4 over 14 weeks and X-rayed periodically using a Faxitron to ensure decalcification. Upon decalcification of the tibia length was measured from all lambs and the proximal epiphyses that included the growth plate was dissected from a pair of sibling lambs (CH2, CH3, affected and unaffected respectively). The tissue was processed for wax embedding using extended wax infiltration steps of 90 minutes. Sections (7 µm thick) were cut and haematoxylin and eosin (H&E) staining was conducted following routine procedures using an automatic stainer (Autostainer XL, Leica Biosystems). Stained slides were scanned on a Hamamatsu SlideScanner and viewed on NanoZoomer Digital Pathology. Measurements were made of the total growth plate height, the height of the hypertrophic zone (HZ), proliferative zone (PZ) and resting zone (RZ), using standard histological features of cell morphology^50,51^.

### Immunofluorescence

Lamb trachea were dissected in PBS, fixed in 4% PFA/PBS overnight before washing in graded sucrose:PBS solutions, embedded in sucrose gelatin blocks and snap frozen on -80°C isopentane. 10 µM sections were cut on a cryostat, mounted on polysine slides and airdried at room temperature for 30 minutes. Sections were washed in 37°C PBS for 30 minutes, blocked in 5% goat serum/0.5% Triton/PBS and incubated with either 1:100 Prickle antibody (proteintech 22589-1-AP)/5% goat serum/0.5% Triton/PBS or 1:100 Prickle antibody (proteintech 22589-1- AP)/5% goat serum/0.5% that had been incubated with 1:500 Prickle fusion protein (proteintech Ag18407) for 45 minutes or in 5% goat serum/0.5% Triton/PBS alone with no Prickle antibody, overnight at 4°C. Slides were subsequently washed in PBS at room temperature for 3 hours, incubated in secondary anti-rabbit antibody (Invitrogen A21207 Alexa Fluor 594; 1:500 5% goat serum/0.5% Triton/PBS) washed at room temperature for 1 hour, incubated with Phalloidin 488 (A12379) was diluted 1:50 with blocking buffer and added to all slides for 30 mins at room temperature, wash in PBS for 30 mins at room temperature, incubated with DAPI diluted at 1:100 for 10 mins, before a further 30 minute wash at room temperature and mounting with ProLong Gold. Sections were imaged on a Zeiss LSM710, with laser levels standardised on control wildtype sections which had be incubated with 1:100 Prickle1 antibody.

### Hybridisation chain reaction (HCR) RNA fluorescence in situ hybridisation

HCR was performed using the Molecular Instruments HCR RNA-FISH v3.0 kit (Molecular Instruments). The protocol followed was modified from the manufacturer’s protocol for whole-mount chicken embryos^37,52^. Split initiator probes (v3.0) for PRICKLE1 (XM_015282428), PTCH1 (NM_204960.2) and SOX9 and SOX9_add (NM_204281.1) were designed by Molecular Instruments.

## Acknowledgements

This study was supported by the Biotechnology and Biological Sciences Research Council through Institute Strategic Programme Grants (BBS/E/D/05191132, BBS/E/D/20211553, BBS/E/D/10002071) to OM, MGD, HAF, CF, DH, LM, VR, JS, RPW, ZHL, SCB, JAW, ALA. The project was also supported a grant (RIDGENE) to JAW, SCB and ALA from the Genesis Faraday Partnership.

