## Supplementary_materials for "A frameshift mutation in *PRICKLE1* causes skeletal dysplasia in Cheviot sheep"

**Supplementary material**

**Supplementary Table 1:** Biosample and sequence accession numbers for whole genome sequence data

**Supplementary Table 2:** Animals sampled for gross anatomy, CT scans, DNA testing

**Supplementary Table 3a:** Gross measurements of tibiae

**Supplementary Table 3b:** Least square means (mm) and p value comparing Tibia length and width in heterozygous lambs (n=9 lambs) and cases (n=9 lambs)

**Supplementary Table 4a:** Descriptive Statistics for Haematoxylin and eosin-stained sections of the tibia growth plates from 1-day-old case (stumpy, CH2) and its 'normal' sibling (CH3) lambs

**Supplementary Table 4b:** Comparison of least square means for Haematoxylin and eosin-stained sections of the tibia growth plates from 1-day-old case (stumpy, CH2) and its 'normal' sibling (CH3) lambs

### Supplementary Figures

**A.**

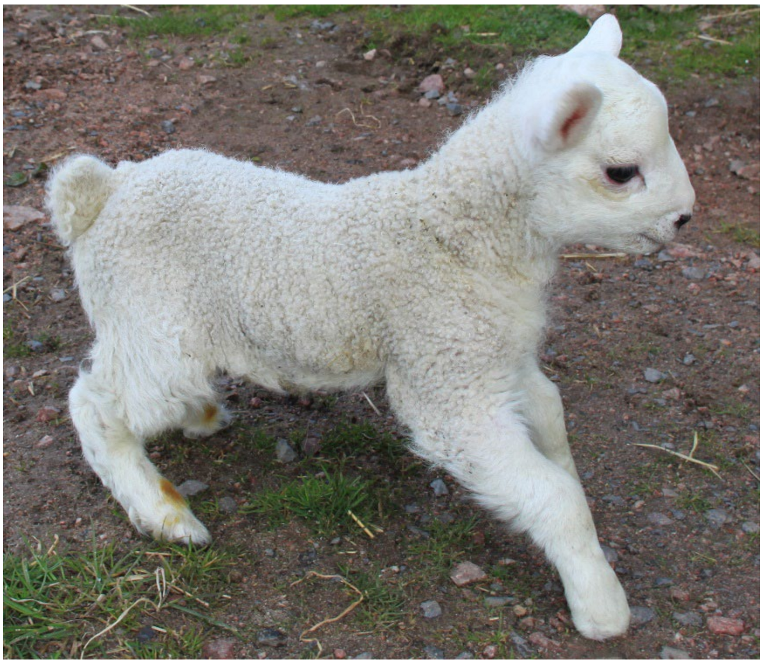

**B.**

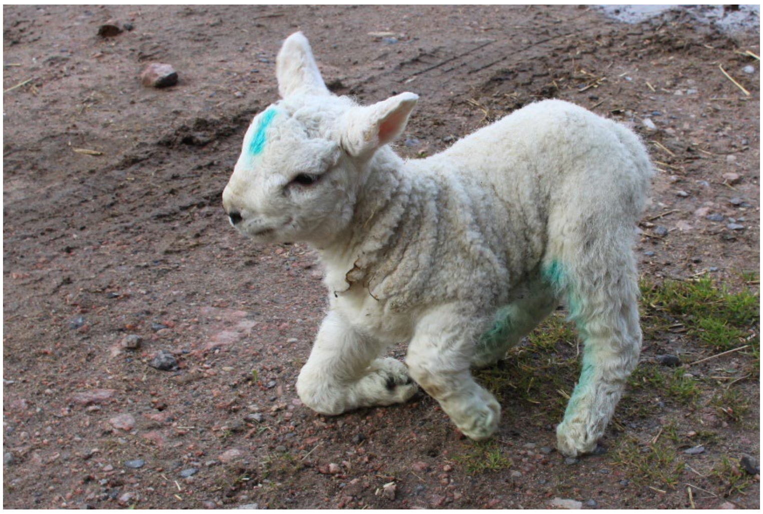

**Supplementary Figure 1. Affected lambs**

A. Affected lamb with domed head, shorter ears, snout and tail evident.
B. Affected lamb with abnormal stance resting on forelimb carpus joint.

**A.**

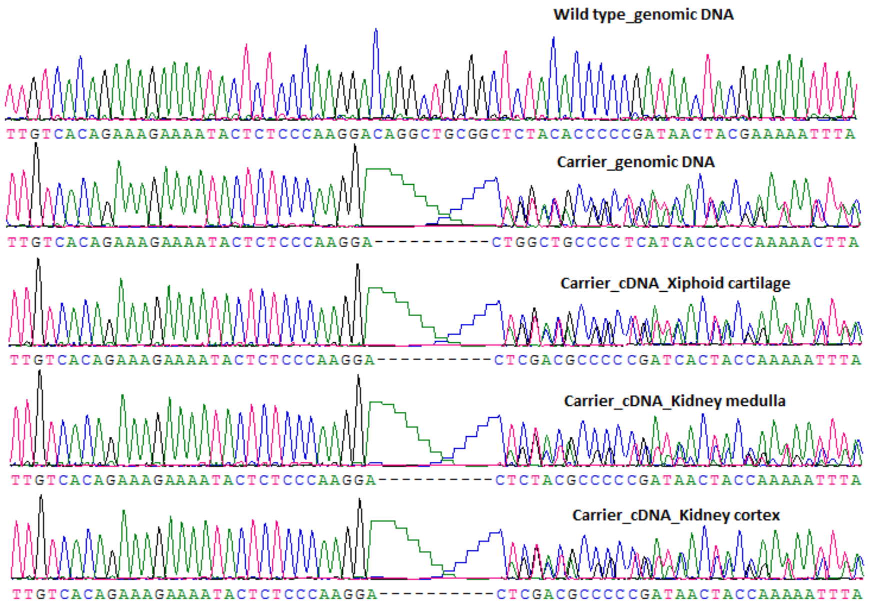


**B.**

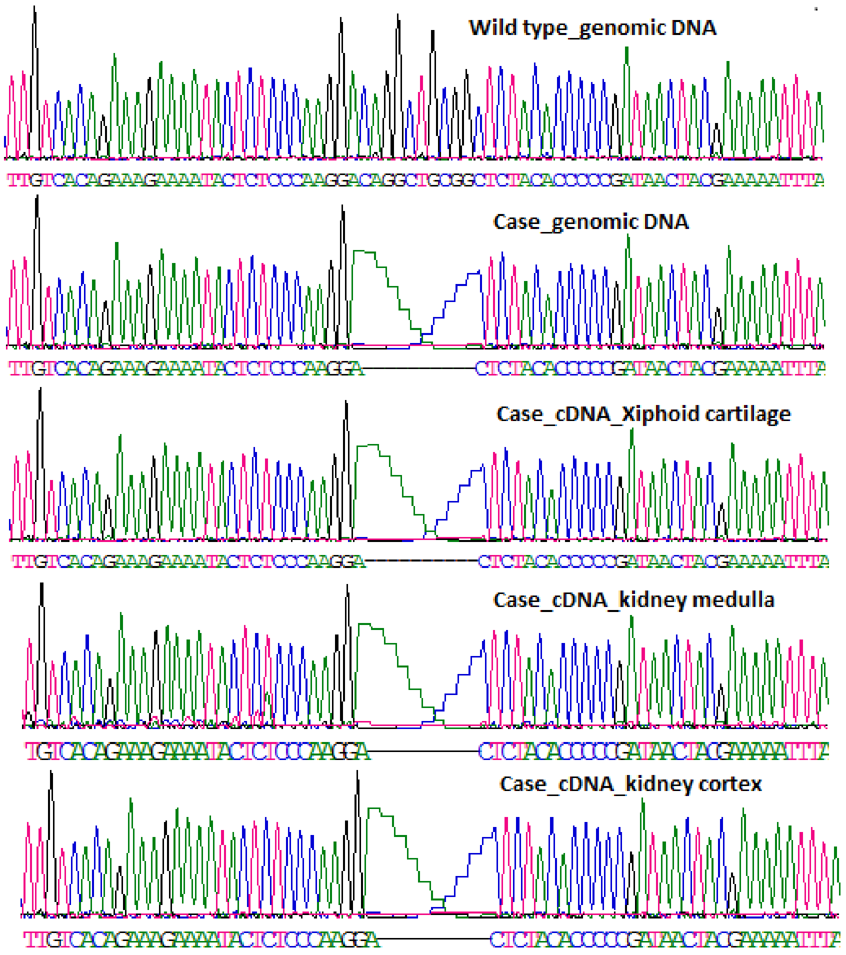


**Supplementary Figure 2: Expression of wild-type and mutant PRICKLE1 alleles. A.** Snapshot of sequence traces of PCR and RT-PCR products obtained using PR1 nested primers to amplify a 688 bp region over the *PRICKLE1* 10 bp deletion. Cheviot CH10 carrier genomic DNA and cDNA from 3 tissues from the same animal are aligned with wild type sequence. The mixed sequence trace observed beyond the deletion indicates the expression of both WT and 10 bp deletion alleles in the carriers, the trace from the carrier genomic DNA shows the same pattern.

**B**. Snapshot of sequence traces of PCR products obtained using PR1 nested primers to amplify a 688 bp region over the *PRICKLE1* 10 bp deletion. Cheviot CH1 case genomic DNA and cDNA from 3 tissues from the same animal are aligned with wild type sequence. The cDNA sequence traces indicate the expression of the *PRICKLE1* 10 bp deletion allele in the cases.

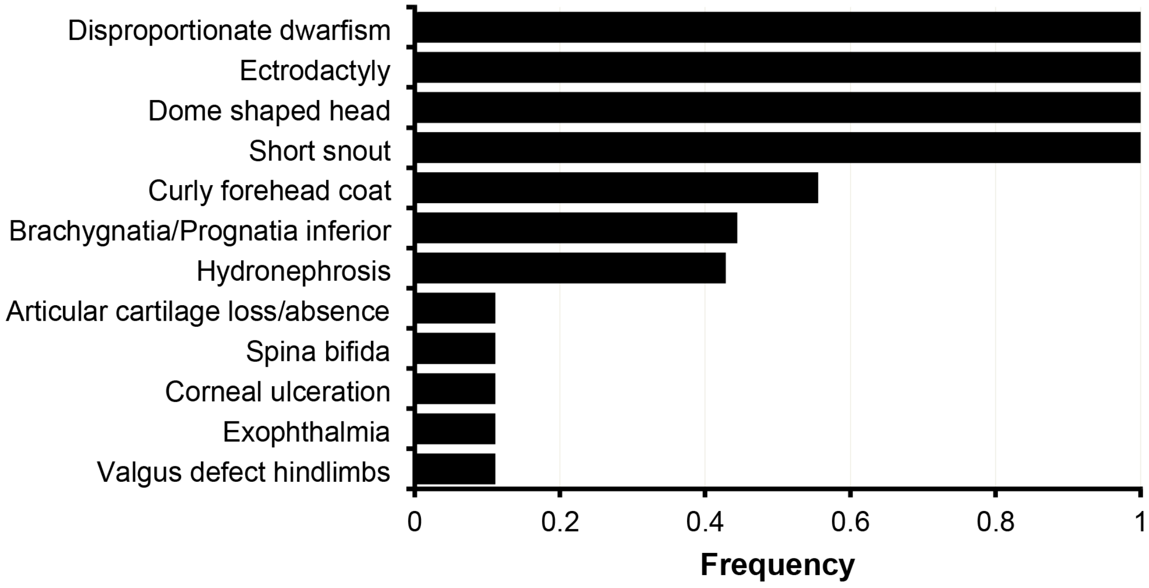


**Supplementary Figure 3: Summary of gross anatomical defects observed in affected lambs at post-mortems.**

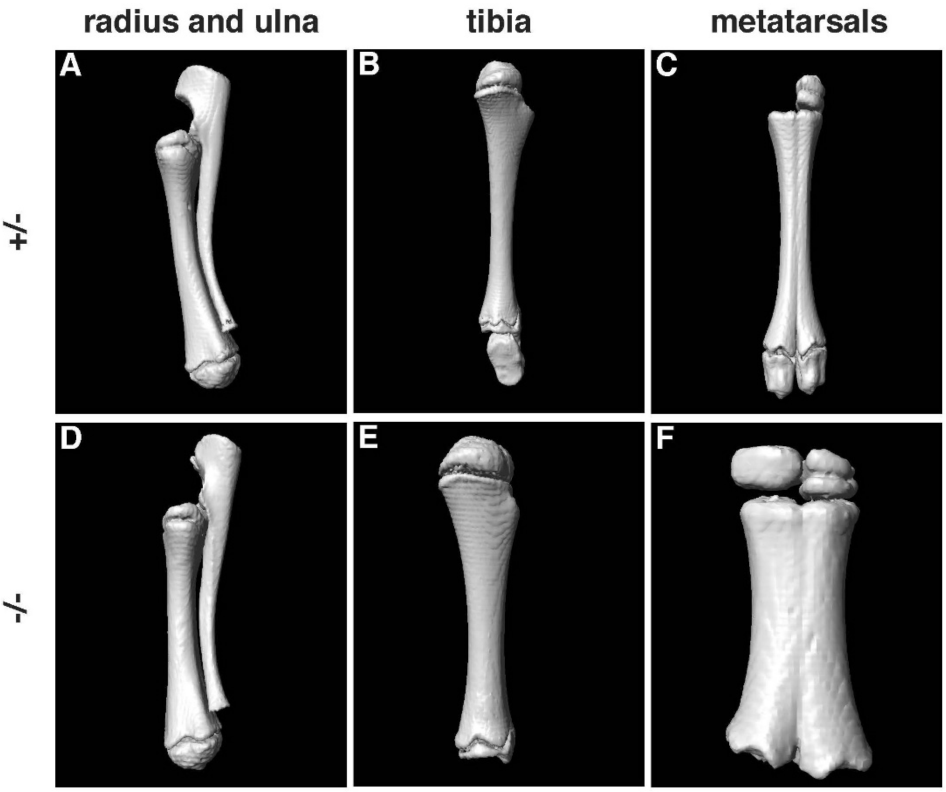


**Supplementary Figure 4: 3D reconstructions of key appendicular bones from *PRICKLE* mutation carriers (A-C) and cases (D-F).** In cases, tibia and metatarsal bones appeared shorter and thicker than those of cases. Note that the missing growth plates in some panels (*e.g.*, C, F) are reconstruction artefacts. Images provided are not to scale with respect to each other.

### Supplementary Tables

**Supplementary Table 1:** Biosample and sequence accession numbers for whole genome sequence data

| name in BioSamples record | BioSamples accession number | Sex | status | ENA sequence accession |
|---|---|---|---|---|
| UK_CHE_F_0000MN237889 | SAMEA103887386 | female | carrier | |
| UK_CHE_F_0000MN237891 | SAMEA103887372 | female | carrier | |
| UK_CHE_F_0000MN237893 | SAMEA103887374 | female | carrier | |
| UK_CHE_F_0000MN237894 | SAMEA103887383 | female | carrier | |
| UK_CHE_M_0000MN237896 | SAMEA103887380 | male | carrier | |
| UK_CHE_M_000000003771 | SAMEA103887385 | male | affected | |
| UK_CHE_M_000000056952 | SAMEA103887371 | male | affected | |
| UK_CHE_F_000000056971 | SAMEA103887375 | female | affected | |
| UK_CHE_M_000000057002 | SAMEA103887381 | male | affected | |
| UK_CHE_M_00062816692B | SAMEA103887389 | male | affected | |

**Supplementary Table 2:** Animals sampled for gross anatomy, CT scans, DNA testing

| Lab ID | Status | Relationships | Sex | Developmental stage | Farm | CT scan | PRICKLE1 DNA result* |
|---|---|---|---|---|---|---|---|
| **CH1** | affected | | female | lamb | S | yes | -/- |
| **CH2** | affected | twin to CH3 | male | lamb | S | yes | -/- |
| **CH3** | unaffected | twin to CH2 | female | lamb | S | yes | +/- |
| **CH4** | putative carrier | mother of CH1 | female | adult | S | | +/- |
| **CH5** | putative carrier | mother of CH2, CH3 | female | adult | S | | +/- |
| **CH6** | affected | | male | lamb | Y | yes | -/- |
| **CH7** | putative carrier | | female | adult | D | | +/- |
| **CH8** | putative carrier | | female | adult | D | | +/- |
| **CH9** | putative carrier | | male | adult | D | | failed |
| **CH10** | unaffected | | male | lamb | S | yes | +/- |
| **CH11** | unaffected | | female | lamb | S | yes | +/- |
| **CH12** | affected | | male | lamb | S | yes | -/- |
| **CH13** | affected | | male | lamb | D | yes | -/- |
| **CH14** | affected | | male | lamb | D | yes | -/- |
| **CH15** | affected | | male | lamb | D | yes | -/- |
| **CH16** | affected | | female | lamb | D | yes | -/- |
| **CH17** | affected | | female | lamb | D | yes | -/- |
| **CH18** | affected | | unknown | lamb | B | | -/- |
| **CH19** | affected | | unknown | lamb | B | | -/- |

* -/- homozygous for deletion in PRICKLE1, +/- heterozygous for deletion in PRICKLE 1

**Supplementary Table 3a:** Gross measurements of tibiae

| Lamb ID | Age (days) | Relationships | Sex | PRICKLE1 DNA result* | Tibia length (mm) | Tibia width (mm) |
|---|---|---|---|---|---|---|
| **CH1** | 3 | | female | -/- | 95 | 18 |
| **CH2** | 1 | twin to CH3 | male | -/- | 90 | 13 |
| **CH3** | 1 | twin to CH2 | female | +/- | 110 | 10 |
| **CH6** | 1 | | female | -/- | 75 | 14 |
| **CH10** | 3 | | male | +/- | 100 | 11 |
| **CH11** | 3 | | female | +/- | 108 | 12 |
| **CH12** | 1 | | male | -/- | 88 | 14 |
| **CH13** | 2 | | male | -/- | 100 | 14 |
| **CH14** | 4 | | male | -/- | 86 | 13 |

* -/- homozygous for deletion in PRICKLE1, +/- heterozygous for deletion in PRICKLE 1

**Supplementary Table 3b:** Least square means (mm) and p value comparing Tibia length and width in heterozygous lambs (n=9 lambs) and cases (n=9 lambs)

| status | Normal | Case | p-value |
|---|---|---|---|
| Tibia_length | 106±4.47 | 89±3.16 | 0.0172 |
| Tibia width | 11±0.96 | 14.3±0.68 | 0.0252 |

**Supplementary Table 4a:** Descriptive Statistics for Haematoxylin and eosin-stained sections of the tibia growth plates from 1-day-old case (stumpy, CH2) and its ‘normal’ sibling (CH3) lambs

| Region | N | mean | se | STD | CV | min | max |
|---|---|---|---|---|---|---|---|
| **HZ** | 105 | 228.56 | 4.66 | 47.7 | 20.87 | 146.12 | 452.09 |
| **PZ** | 103 | 336.72 | 8.49 | 86.21 | 25.6 | 196.08 | 592.68 |
| **RZ** | 104 | 526.73 | 12.69 | 129.19 | 24.53 | 337.60 | 1220.67 |
| **Total** | 100 | 1085.54 | 15.29 | 152.9 | 14.08 | 791.35 | 1477.57 |

HZ = hypertrophic zone, PZ = proliferative zone, RZ = resting zone (see Figure 6)

**Supplementary Table 4b:** Comparison of least square means for Haematoxylin and eosin-stained sections of the tibia growth plates from 1-day-old case (stumpy, CH2) and its ‘normal’ sibling (CH3) lambs

| Region | Mean | | | | R2 | P_value |
|---|---|---|---|---|---|---|
| | Lamb1 (CH2) | se | Lamb2 (CH3) | se | | |
| **HZ** | 222.33 | 6.79 | 234.01 | 6.36 | 0.02 | 0.212 |
| **PZ** | 256.46 | 6.92 | 401.49 | 6.22 | 0.71 | <.0001 |
| **RZ** | 510.99 | 18.62 | 540.22 | 17.24 | 0.01 | 0.2519 |
| **Total** | 994.12 | 19.23 | 1160.34 | 17.39 | 0.30 | <.0001 |
